## Supplementary materials for "Non-rapid eye movement sleep and wake neurophysiology in schizophrenia"

### *Supplementary table 1: Group differences in EEG parameters between SCZ and CTR*

| Sleep microstructure metrics | SCZ Mean ± SD(min, max) | CTR Mean ± SD(min, max) | Number of channels with $p_{adj}$<0.05 | Significant $p_{adj}$ range | Effect size across channels with $p_{adj}$<0.05 |
|---|---|---|---|---|---|
| ***SS density, n/min*** | 1.3±0.49 (0.7–2.1) | 1.5±0.47 (0.9-2.2) | 20 | 3e-04 - 0.0117 | -0.79 - -0.6 |
| ***SS amplitude, uV*** | 24.4±6.73 (16.9–33.3) | 28.6±6.73 (20-37.5) | 49 | 3e-04 - 0.0467 | -0.82 - -0.43 |
| ***SS ISA, a.u.*** | 1.3±0.24 (1.2–1.5) | 1.5±0.2 (1.3-1.7) | 34 | 3e-04 - 0.048 | -0.96 - -0.63 |
| SS duration, s | 0.9±0.09 (0.8–0.9) | 0.9±0.07 (0.8-0.9) | 0 | | |
| SS chirp, a.u. | -0.23±0.151 (-0.31–-0.14) | -0.25±0.117 (-0.34--0.15) | 0 | | |
| SS frequency, Hz | 11.1±0.41 (10.9–11.2) | 11.2±0.48 (11-11.4) | 0 | | |
| **FS density, n/min** | 1.7±0.82 (1–2.6) | 2.4±0.8 (1.4-3.3) | 53 | 3e-04 - 0.0443 | -1.27 - -0.56 |
| **FS amplitude, uV** | 17.2±4.62 (11.8–23) | 19.5±5.3 (12.4-27.1) | 22 | 0.0013 - 0.0457 | -0.62 - -0.41 |
| **FS ISA, a.u.** | 1.3±0.18 (1.2–1.4) | 1.4±0.15 (1.3-1.5) | 1 | 0.0373 | -0.73 |
| **FS duration, s** | 0.8±0.09 (0.7–0.8) | 0.8±0.07 (0.8-0.9) | 35 | 0.0057 - 0.0477 | -0.98 - -0.69 |
| **FS chirp, a.u.** | -0.22±0.109 (-0.27–-0.16) | -0.17±0.101 (-0.25--0.09) | 20 | 0.0027 - 0.0437 | -1.09 - -0.59 |
| FS frequency, Hz | 13.8±0.37 (13.6–13.9) | 13.7±0.37 (13.4-13.8) | 0 | | |
| **SO density, n/min** | 13.3±2.16 (11.9–14.6) | 12.3±1.68 (10.1-13.7) | 30 | 3e-04 - 0.0483 | 0.59 - 1.16 |
| **SO duration, s** | 1.1±0.22 (0.9–1.3) | 1±0.14 (0.8-1.1) | 36 | 3e-04 - 0.033 | 0.66 - 1.92 |
| **SO slope, a.u.** | 171.1±42.04 (107.9–249.2) | 208.3±48.59 (122.7-323.2) | 44 | 3e-04 - 0.0377 | -1.14 - -0.55 |
| SO neg peak amplitude, uV | 36.2±8.37 (25.5–49.2) | 38.5±8.7 (25-54.4) | 0 | | |
| SO peak-to-peak amplitude, uV | 61.3±13.96 (43.9–80.9) | 64±13.8 (42.2-88.9) | 0 | | |
| SS SO coupling strength, z | 2.4±2.37 (0.8–4.5) | 2.8±2.34 (1.2-4.6) | 0 | | |
| **SS SO coupling overlap, z** | 2.1±1.08 (1.2–2.8) | 2.7±0.91 (1.7-3.4) | 13 | 0.0063 - 0.046 | -1.14 - -0.62 |
| **SO phase angle when SS occur,°** | 352.5±70.01 (326.7–374.9) | 373.8±56.91 (354.5-389.9) | 2 | 3e-04 - 0.019 | -1.07 - -1 |
| FS SO coupling strength, z | 6.7±4.06 (3.8–9.5) | 5.8±2.97 (3.6-8) | 0 | | |
| FS SO coupling overlap, z | 2.9±1.26 (2.2–3.4) | 2.9±1.03 (1.9-3.5) | 0 | | |
| **SO phase angle when FS occur,°** | 233±22.52 (218.8–241.7) | 240.8±26.8 (220.6-252.2) | 2 | 0.045 - 0.0497 | -0.65 - -0.41 |
| **MMN Amplitude, uV** | 1.3±0.98 (2–0.6) | 1.6±0.99 (2.6-0.6) | 1 | 0.012 | -0.7 |
| MMN Latency, s | 180.7±25.65 (179–183.8) | 182.3±26.65 (178.7-186.2) | 0 | | |
| **P50 S2/S1 ratio** | 0.8±0.4 (0.7–1) | 0.6±0.39 (0.4-0.8) | 10 | 0.001 - 0.044 | 0.62 - 1.06 |
| P50 S1 amplitude, uV | 2.1±1.04 (1.6–2.9) | 2.2±1.04 (1.5-3.1) | 0 | | |
| P50 S2 amplitude, uV | 1.5±0.82 (1.2–1.8) | 1.2±0.72 (0.9-1.5) | 0 | | |
| **ASSR Power, dB** | -0.7±0.42 (-1–-0.4) | -0.5±0.43 (-0.8--0.2) | 15 | 0.0077 - 0.048 | -0.69 - -0.56 |
| **ASSR Phase synchrony, a.u.** | 0.2±0.1 (0.2–0.3) | 0.3±0.13 (0.2-0.4) | 20 | 0.0013 - 0.046 | -0.73 - -0.54 |

***Supplementary table 2: prescribed medications in the SCZ sample***

| Medication | Number of subjects (total 72*) |
|---|---|
| ***Antipsychotics*** | 69 |
| Amisulpride | 22 |
| Chlorpromazine Hydrochloride | 1 |
| Aripiprazole | 12 |
| Olanzapine | 26 |
| Clozapine | 12 |
| Quetiapine Fumarate | 6 |
| Risperidone | 14 |
| Paliperidone (sustained release) | 1 |
| ***Emotion stabilizer and Antiepileptic*** | 17 |
| ***Anticholinergics*** | 10 |
| ***Sedatives and Tranquilizers*** | 13 |
| ***Antidepressants*** | 4 |

*- due to 4 subjects being removed there were 67 out of 68. The table above shows the full dataset

### *Supplementary table 3: Medication associations with EEG metrics within the SCZ sample*

| EEG metric | CPZ equivalent antipsychotic dose (n=67) | Antipsychotic medication | | | | | | Adjunctive medication | | |
|---|---|---|---|---|---|---|---|---|---|---|
| | | Amisulpride (n=22) | Aripiprazole (n=12) | Olanzapine (n=26) | Clozapine (n=12) | Quetiapine Fumarate (n=6) | Risperidone (n=14) | Sedatives and tranquilizers (n=13) | Mood stabilizers and antiepileptics (n=17) | Anticholinergics (n=10) |
| SS Density (↓ in SCZ) | | | | ↑ FPZ,FP1,FP2 | ↓ AF3,F7,T7 | | | | | |
| FS Density (↓ in SCZ) | | | | | ↓ 9 channels | | | | | |
| SS Amplitude (↓ in SCZ) | | | | | ↓ AFZ,T7 | | | | | ↑ P2,P4 |
| FS Amplitude (↓ in SCZ) | | | | | | | | | ↓ AFZ | |
| SS ISA (↓ in SCZ) | | | ↑ F2,F4 | | ↓ T7 | | | | | |
| FS ISA (↓ in SCZ) | | | | | | | | | | |
| FS Duration (↓ in SCZ) | | | | | ↓ 7 channels | | | | | |
| FS Chirp (↓ in SCZ) | ↑FC6, ↓TP8 | | | | ↓ 8 channels | ↓ FC1 | | | | |
| SO Density (↑ in SCZ) | | | | | | | | | | |
| SO Duration (↑ in SCZ) | | | | | | | | ↑ 10 channels | | |
| SO Slope (↓ in SCZ) | | | | | | | | ↓ 8 channels | ↓ 23 channels | |
| SS overlap with SO (↓ in SCZ) | | | ↑ 6 channels | | ↑ CP5,CZ,F5 | | | | ↓ 6 channels | |
| SO phase angle when SS occur (↓ in SCZ) | | | ↑ POZ | | | | | | ↓ CPZ,CZ | ↑ C3,C4,C5 |
| SO phase angle when FS occur (↓ in SCZ) | | | | | | | | | | ↑ CP3 |
| PSD PC #4 (↓ in SCZ) | | | | | ↓ | | | | | |
| PSI PC #1 (↑ in SCZ) | | | | | | | | | | |
| MMN Amplitude (↓ in SCZ) | | | ↑ AF4 | | | | | | | |
| P50 S2/S1 ratio (↑ in SCZ) | | | | | | | | | | |
| ASSR Power (↓ in SCZ) | | | | | | | | | | |
| ASSR Phase synchrony (↓ in SCZ) | | | | ↑ 5 channels | | | | | | ↓ F8 |

*A linear regression model with formula EEG metric ~ medication + sex + age + error was fit for each channel for each EEG metric, where medication was a vector with continuous data for Chlorpromazine (CPZ) equivalent antipsychotic dose and a binary vector (True or False for medication use) for other columns. Channels where the association was significant (unadjusted $p<0.01$) are provided in the table together with the direction of effect (↓ or ↑ in an EEG metric with medication)*

*Supplementary table 4: Significance of group differences between SCZ and CTR adjusted for medication*

| **EEG metric** | **Antipsychotic medication** | | | | | | **Adjunctive medication** | | |
|---|---|---|---|---|---|---|---|---|---|
| | **Amisulpride (n=22)** | **Aripiprazole (n=12)** | **Olanzapine (n=26)** | **Clozapine (n=12)** | **Quetiapine Fumarate (n=6)** | **Risperidone (n=14)** | **Sedatives and tranquilizers (n=13)** | **Emotion stabilizers and antiepileptics (n=17)** | **Anticholinergics (n=10)** |
| SS Density, 20 channel(s) | | | | | | | | | |
| FS Density, 53 channel(s) | | | | | | | | | |
| SS Amplitude, 49 channel(s) | | | | | | | | | |
| FS Amplitude, 22 channel(s) | | | | | | | | | |
| SS ISA, 34 channel(s) | | | | | | | | | |
| **FS ISA, 1 channel(s)** | n.s. | | n.s. | | | | | n.s. | |
| FS Duration, 35 channel(s) | | | | | | | | | |
| FS Chirp, 20 channel(s) | | | | | | | | | |
| SO Density, 30 channel(s) | | | | | | | | | |
| SO Duration, 36 channel(s) | | | | | | | | | |
| SO Slope, 44 channel(s) | | | | | | | | | |
| SS overlap with SO, 13 channel(s) | | | | | | | | | |
| SO phase angle when SS occur, 2 channel(s) | | | | | | | | | |
| **SO phase angle when FS occur, 2 channel(s)** | | | n.s. | | | | | n.s. | |
| PSD PC #4 | | | | | | | | | |
| PSI PC #1 | | | | | | | | | |
| **MMN Amplitude, 1 channel(s)** | | | n.s. | | | | | | |
| P50 S2/S1 ratio, 10 channel(s) | | | | | | | | | |
| ASSR Power, 15 channel(s) | | | | | | | | | |
| ASSR Phase synchrony, 20 channel(s) | | | | | | | | | |

*For each medication specified in the columns, SCZ patients taking that medication were excluded from the SCZ sample and group differences for each EEG metric at each channel were re-estimated.* ***n.s.*** *indicates that there were no channels with significant differences (unadjusted p<0.01) once subjects taking the corresponding medication were excluded.*

*Supplementary table 5 Association between clinical variables and EEG metrics in the SCZ sample*

| EEG metric | SCZ Duration | PANSS 5 factors | | | | |
|---|---|---|---|---|---|---|
| | | Positive | Negative | Disorganized Concrete | Excited | Depressed |
| SS Density (↓ in SCZ) | | | ↓ AF3,AF4,F8 | ↓ 24 channels | | |
| FS Density (↓ in SCZ) | | | | | | |
| SS Amplitude (↓ in SCZ) | | | ↓ 15 channels | | | |
| FS Amplitude (↓ in SCZ) | | | ↓ 5 channels | | | |
| SS ISA (↓ in SCZ) | | | | | | |
| FS ISA (↓ in SCZ) | | ↑ F6,F7,F8,FC6 | | | | |
| FS Duration (↓ in SCZ) | | | | | | |
| FS Chirp (↓ in SCZ) | | | | | | |
| SO Density (↑ in SCZ) | ↓ 17 channels | | | | | |
| SO Duration (↑ in SCZ) | | | ↓ F6 | | | |
| SO Slope (↓ in SCZ) | | | | | ↑ FT8,T8,TP7 | |
| SS overlap with SO (↓ in SCZ) | | | | | | |
| SO phase angle when SS occur (↓ in SCZ) | ↑ FPZ,FP2 | | | ↓ FCZ | ↓ FC2,FCZ | |
| SO phase angle when FS occur (↓ in SCZ) | ↓ FT8,O1,P8,TP8 | | | | | |
| PSD PC #4 (↓ in SCZ) | | | | | | |
| PSI PC #1 (↑ in SCZ) | | | | | | |
| MMN Amplitude (↓ in SCZ) | | | ↓ 7 channels | ↓ FT7,T7 | | |
| P50 S2/S1 ratio (↑ in SCZ) | | ↑ FZ | ↑ C1,CZ | | ↑ F1 | |
| ASSR Power (↓ in SCZ) | ↓ O1 | | | | | |
| ASSR Phase synchrony (↓ in SCZ) | | | | | | |

*A linear regression model with formula EEG metric ~ clinical variable + sex + age + error was fit for each channel / EEG metric. Channels with significant association (unadjusted p<0.01) are listed together with the direction of effect (↓ or ↑ in an EEG metric with clinical variable)*

*Supplementary table 6. Demographic characteristics of the independent samples*

| | **Lunesta dataset** (4-7 EEG channels: F3, F4, C3,C4,Pz,O1,O2) | | **ESZ dataset** (58 EEG channels) | | **GCRC dataset** (4 EEG channels: C3,C4,O1,O2) | |
|---|---|---|---|---|---|---|
| **Sample characteristics** | **SCZ** | **CTR** | **SCZ** | **CTR** | **SCZ** | **CTR** |
| N | 20 | 17 | 26 | 29 | 11 | 13 |
| Sex | 5 females | 3 females | 5 females | 8 females | 3 females | 3 females |
| Race | White - 15; Black - 3; Asian - 1; | White - 17 | White - 12; Black - 6; Asian - 3; na - 5 | White - 17; Asian - 3; na - 9 | White - 9; Black - 1; Native American - 1; | White - 11; Black - 1; Multiracial - 1; |
| Age, years | 34.9±8.69 | 36.3±7.12 | 32.3±7.53 | 30.1±6.25 | 44.1±9.5 | 43.1±6.11 |
| Parental education, years | 15.1±3.33 | 13.5±2.15 | 13.4±3.01 | **16.7±2.02*** | 13.1±1.73 | 13.5±3.02 |
| Std. antipsychotic dose, mg | 314±240.15 | | 416.5±310 | | 567.2±512.87 | |
| **Sleep macrostructure parameters** | | | | | | |
| Total time in bed, mins | **622±75*** | 578±44 | 575±42.2 | 561±42.5 | **610±70*** | 563±50.1 |
| Total sleep time, mins | **464±72.6*** | 420±52.4 | 510±71.1 | 506±45.7 | 430±101.7 | 390±71 |
| Sleep latency, mins | **19±13.7*** | 8±4.9 | 26±17.5 | 15±10.2 | 106±66.7 | 113±30.3 |
| Wake after sleep onset, mins | 33±33.1 | 35±17.5 | **22±16.8*** | 33±19.2 | 62±55.7 | 47±23.3 |
| Sleep efficiency (/TIB) | 87±8.6 | 88±4.9 | 90±6.6 | 91±3.6 | 71±14.2 | 69±8 |
| Sleep efficiency (/start-end of sleep) | 92±8.6 | 92±3.4 | **96±3*** | 94±3.5 | 86±10.9 | 88±5.7 |
| Duration N1, mins (%) | 33±22.7 (8) | 37±16.1 (9) | 44±22.4 (9) | 52±22 (10) | 32±18.5 (8) | 24±13.3 (6) |
| Duration N2, mins (%) | 233±68.7 (53) | 234±36.5 (56) | 266±79 (53) | 265±54.4 (52) | 297±94.6 (69) | 240±66.6 (64) |
| Duration N3, mins (%) | 83±50.4 (18) | 56±23 (13) | 93±55.3 (19) | 93±32 (19) | 44±39.5 (10) | 42±20.8 (11) |
| Duration REM, mins (%) | 97±38.2 (22) | 92±22 (22) | 88±29.7 (17) | 92±29.6 (18) | 57±31.1 (13) | 84±42.9 (21) |
| Latency of REM, mins | 150±84.9 | 100±49.1 | 123±59.6 | 106±40.2 | **195±72.6*** | 114±76.4 |
| Number of cycles | 4±1.2 | 5±1.2 | 5±1.4 | 5±1 | 4±1.2 | 4±1.2 |
| Cycle length, mins | **116±34.4*** | 94±19.4 | 112±23.4 | 97±15.1 | 115±38.7 | 96±21.1 |

* *p-value<0.05*

*Supplementary table 7 Replication analysis for spindle, slow oscillation and coupling metrics*

| Sleep microstructure metrics | EEG channel | GRINS | | | | Replication | | | |
|---|---|---|---|---|---|---|---|---|---|
| | | SCZ Mean ± SD | CTR Mean ± SD | p-value | Effect size., SD | SCZ Mean ± SD | CTR Mean ± SD | p-value | Effect size SD |
| **SS density, n/min** | C3 | 1.2±0.45 | 1.4±0.4 | **0.0179** | -0.45 | 1±0.43 | 1.2±0.47 | **0.0053** | -0.51 |
| | C4 | 1.1±0.47 | 1.3±0.42 | **0.0312** | -0.45 | 0.9±0.43 | 1.2±0.47 | **0.004** | -0.58 |
| | O1 | 0.7±0.35 | 0.9±0.35 | **7E-04** | -0.66 | 0.6±0.3 | 0.7±0.33 | **0.0139** | -0.46 |
| | O2 | 0.8±0.38 | 1±0.32 | **0.0079** | -0.59 | 0.5±0.28 | 0.7±0.31 | **0.0012** | -0.61 |
| **FS density, n/min** | C3 | 1.9±0.9 | 2.7±0.78 | **0** | -1.03 | 1.7±0.94 | 2.6±0.76 | **0** | -1.14 |
| | C4 | 2±0.86 | 2.8±0.75 | **0** | -1.14 | 1.6±0.93 | 2.3±0.84 | **0** | -0.91 |
| | O1 | 1.8±0.99 | 2.4±1.04 | **0.0011** | -0.59 | 1.2±0.95 | 1.8±1.02 | **9E-04** | -0.61 |
| | O2 | 1.8±0.98 | 2.5±1.07 | **9E-04** | -0.63 | 1.2±0.97 | 1.8±0.94 | **9E-04** | -0.61 |
| SS amplitude, uV | C3 | 24.9±6.46 | 29.5±5.99 | **0** | -0.78 | 24.1±5.95 | 25.5±7.66 | 0.3713 | -0.19 |
| | C4 | 25.9±6.7 | 31.3±7.22 | **1E-04** | -0.75 | 25±6.6 | 25.4±6.1 | 0.7147 | -0.06 |
| | O1 | 18.2±5.75 | 21.2±6.08 | **0.0064** | -0.49 | 16.9±4.66 | 18±6.59 | 0.4946 | -0.17 |
| | O2 | 18.6±5.78 | 21.8±6.37 | **0.0026** | -0.51 | 17.4±5.08 | 18.3±6.05 | 0.3522 | -0.14 |
| **FS amplitude, uV** | C3 | 18.5±4.59 | 21.6±5.54 | **0.0021** | -0.57 | 16.7±4.42 | 19.6±6.08 | **0.017** | -0.48 |
| | C4 | 19.7±5.14 | 22.6±5.27 | **0.0015** | -0.55 | 17.2±4.62 | 19±4.77 | 0.0782 | -0.37 |
| | O1 | 14.2±4.95 | 15.3±5.26 | 0.1123 | -0.22 | 11.5±3.34 | 13±4.91 | 0.1274 | -0.29 |
| | O2 | 14.5±4.73 | 16.3±5.99 | **0.0389** | -0.3 | 11.7±3.75 | 13.1±4.56 | 0.1051 | -0.31 |
| **SS ISA, a.u.** | C3 | 1.3±0.2 | 1.4±0.2 | **4E-04** | -0.9 | 1.2±0.19 | 1.4±0.17 | **1E-04** | -0.92 |
| | C4 | 1.3±0.22 | 1.4±0.19 | **0.0012** | -0.82 | 1.2±0.2 | 1.3±0.18 | **0** | -0.86 |
| | O1 | 1.3±0.31 | 1.4±0.24 | **0.0303** | -0.65 | 1.2±0.25 | 1.4±0.27 | **1E-04** | -0.72 |
| | O2 | 1.3±0.28 | 1.4±0.19 | **0.0049** | -0.83 | 1.2±0.3 | 1.4±0.28 | **0.0016** | -0.57 |
| **FS ISA, a.u.** | C3 | 1.3±0.18 | 1.4±0.13 | 0.084 | -0.48 | 1.3±0.21 | 1.4±0.15 | **0.0287** | -0.49 |
| | C4 | 1.3±0.18 | 1.4±0.16 | **0.0492** | -0.54 | 1.3±0.21 | 1.4±0.16 | 0.0746 | -0.42 |
| | O1 | 1.3±0.2 | 1.4±0.15 | 0.1626 | -0.37 | 1.2±0.21 | 1.3±0.19 | **0.0027** | -0.58 |
| | O2 | 1.4±0.18 | 1.4±0.15 | 0.0784 | -0.41 | 1.2±0.2 | 1.3±0.18 | **0.0023** | -0.56 |
| **FS duration, s** | C3 | 0.8±0.09 | 0.8±0.07 | **0.0032** | -0.79 | 0.8±0.12 | 0.8±0.09 | **0.0459** | -0.45 |
| | C4 | 0.8±0.09 | 0.8±0.07 | **0.0031** | -0.82 | 0.8±0.11 | 0.8±0.09 | 0.0596 | -0.41 |
| | O1 | 0.8±0.11 | 0.9±0.07 | **0.0056** | -0.85 | 0.8±0.11 | 0.8±0.09 | **0.0134** | -0.62 |
| | O2 | 0.8±0.11 | 0.9±0.08 | **0.0119** | -0.7 | 0.8±0.12 | 0.8±0.09 | **0.0072** | -0.55 |
| **FS chirp, Hz** | C3 | -0.2±0.09 | -0.2±0.1 | 0.1766 | -0.3 | -0.2±0.09 | -0.2±0.1 | 0.1655 | -0.26 |
| | C4 | -0.2±0.1 | -0.2±0.1 | **0.0177** | -0.47 | -0.2±0.1 | -0.2±0.09 | 0.5148 | -0.07 |
| | O1 | -0.2±0.11 | -0.1±0.08 | **6E-04** | -1.09 | -0.1±0.1 | -0.1±0.07 | **0.0379** | -0.51 |

| | | | | | | | | | |
|---|---|---|---|---|---|---|---|---|---|
| | O2 | -0.2±0.1 | -0.1±0.08 | **4E-04** | -0.94 | -0.1±0.08 | -0.1±0.07 | 0.0854 | -0.33 |
| SO density, n/min | C3 | 13.4±2.08 | 12.5±1.59 | **0.0144** | 0.57 | 12.9±1.99 | 12.3±1.28 | 0.3876 | 0.44 |
| | C4 | 13.3±2.1 | 12.5±1.58 | **0.023** | 0.52 | 13±2.07 | 12.1±1.29 | 0.0615 | 0.72 |
| | O1 | 12.1±2.16 | 10.2±1.61 | **0** | 1.16 | 11.2±1.59 | 10.2±1.43 | **0.0033** | 0.67 |
| | O2 | 11.9±2.13 | 10.2±1.63 | **0** | 1.06 | 11±1.64 | 10.2±1.28 | **0.0108** | 0.61 |
| SO duration, s | C3 | 1±0.18 | 0.9±0.12 | **2E-04** | 1.08 | 1±0.12 | 0.9±0.1 | 0.0884 | 0.52 |
| | C4 | 1±0.19 | 0.9±0.09 | **0** | 1.67 | 0.9±0.11 | 0.9±0.12 | 0.2306 | 0.22 |
| | O1 | 1.3±0.32 | 1.1±0.2 | **0.0022** | 0.93 | 1.1±0.22 | 1±0.17 | 0.6103 | 0.32 |
| | O2 | 1.3±0.32 | 1.1±0.18 | **0.0013** | 1.16 | 1.1±0.18 | 1±0.15 | 0.2733 | 0.46 |
| SO slope, a.u. | C3 | 181.2±41.52 | 222.8±45.87 | **0** | -0.91 | 194.7±43.3 | 198.8±38.16 | 0.7308 | -0.11 |
| | C4 | 188±43.82 | 236.7±48.31 | **0** | -1.01 | 194.2±44.34 | 197±33.89 | 0.9467 | -0.08 |
| | O1 | 107.9±32.63 | 122.7±33.92 | **0.0451** | -0.44 | 108.2±28.98 | 108±29.97 | 0.7957 | 0 |
| | O2 | 110.4±31.75 | 134.3±41.45 | **0.0027** | -0.58 | 108.5±29.44 | 110.4±29.13 | 0.999 | -0.07 |
| SS SO coupling overlap, z | C3 | 2.3±0.94 | 2.8±0.91 | **0.0065** | -0.57 | 2.6±1.2 | 2.9±0.73 | 0.2632 | -0.53 |
| | C4 | 2.1±1.05 | 2.8±0.9 | **4E-04** | -0.79 | 2.2±1.61 | 2.9±0.69 | 0.0872 | -1.06 |
| | O1 | 1.3±1.24 | 2±1.06 | **0.0018** | -0.75 | 1.1±1.34 | 2±1.1 | **0.0016** | -0.83 |
| | O2 | 1.3±1.33 | 1.9±1.09 | **0.002** | -0.63 | 1.1±1.39 | 1.9±1.21 | **0.0048** | -0.68 |
| SO phase angle at SS peak,° | C3 | 356±57.8 | 379.7±48.8 | 0.0776 | -0.49 | 386.7±56.66 | 382.7±53.21 | 0.6712 | 0.07 |
| | C4 | 361.9±57.73 | 383.8±47.97 | **0.026** | -0.46 | 380.4±51.89 | 389.1±45.66 | 0.3559 | -0.19 |
| | O1 | 359.1±108.55 | 384.9±85.24 | 0.3365 | -0.3 | 376±88.78 | 385.9±89.2 | 0.7236 | -0.11 |
| | O2 | 338.2±116.09 | 368.5±93.09 | 0.0595 | -0.33 | **360.2±100.05** | **414.1±87.04** | **0.0088** | **-0.62** |
| SO phase angle at FS peak,° | C3 | 239.2±15.03 | 240.3±22.24 | 0.8406 | -0.05 | 267.9±27.54 | 266.6±26.47 | 0.95 | 0.05 |
| | C4 | 237.3±14.54 | 243±17.51 | 0.1556 | -0.32 | 269.3±31.25 | 262.4±33.87 | 0.7349 | 0.2 |
| | O1 | 218.8±26.06 | 225.9±38.68 | 0.2162 | -0.18 | 261.3±29.21 | 253.6±38.16 | 0.6079 | 0.2 |
| | O2 | 222.1±29.36 | 220.6±52.05 | 0.5997 | 0.03 | 257.1±31.71 | 253.3±34.33 | 0.7406 | 0.11 |
| SS phase-frequency coupling, a.u. | C3 | 0.7±0.22 | 0.8±0.17 | 0.0095 | -0.74 | 0.6±0.3 | 0.7±0.2 | 0.0514 | -0.57 |
| | C4 | 0.6±0.26 | 0.8±0.18 | 0.0016 | -0.9 | 0.7±0.21 | 0.7±0.24 | 0.5336 | -0.17 |
| | O1 | 0.5±0.24 | 0.5±0.26 | 0.2754 | -0.21 | 0.4±0.3 | 0.5±0.27 | 0.386 | -0.21 |
| | O2 | 0.5±0.26 | 0.6±0.27 | 0.1129 | -0.4 | 0.4±0.31 | 0.5±0.26 | 0.0643 | -0.46 |

***Supplementary figure 1: Topographical distribution of coupling characteristics averaged across SCZ and CTR groups***

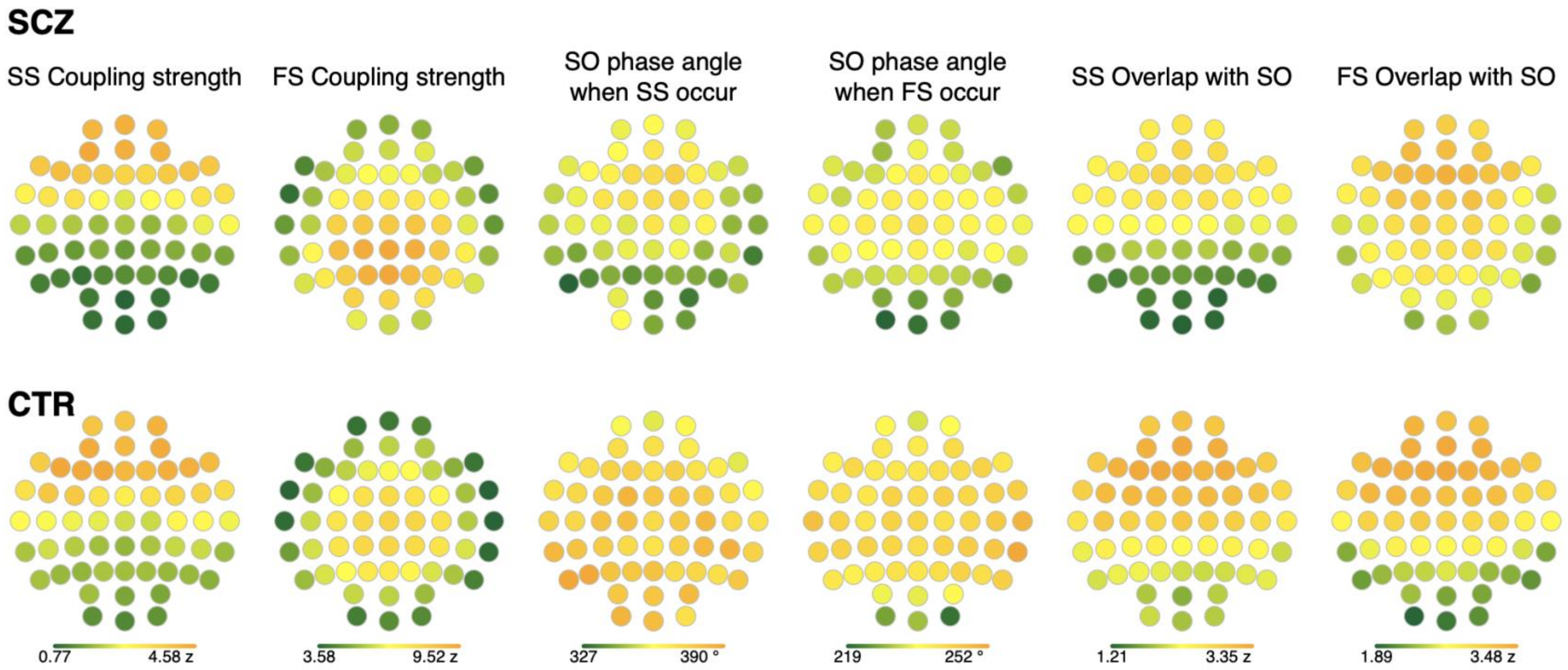


*Topographical distributions of coupling parameters between slow spindles (SS) or fast spindles (FS) and slow oscillations (SO) averaged across SCZ (first row) and CTR (second row).*

*Supplementary figure 2: Instanteneous frequency and its coupling with SO phase*

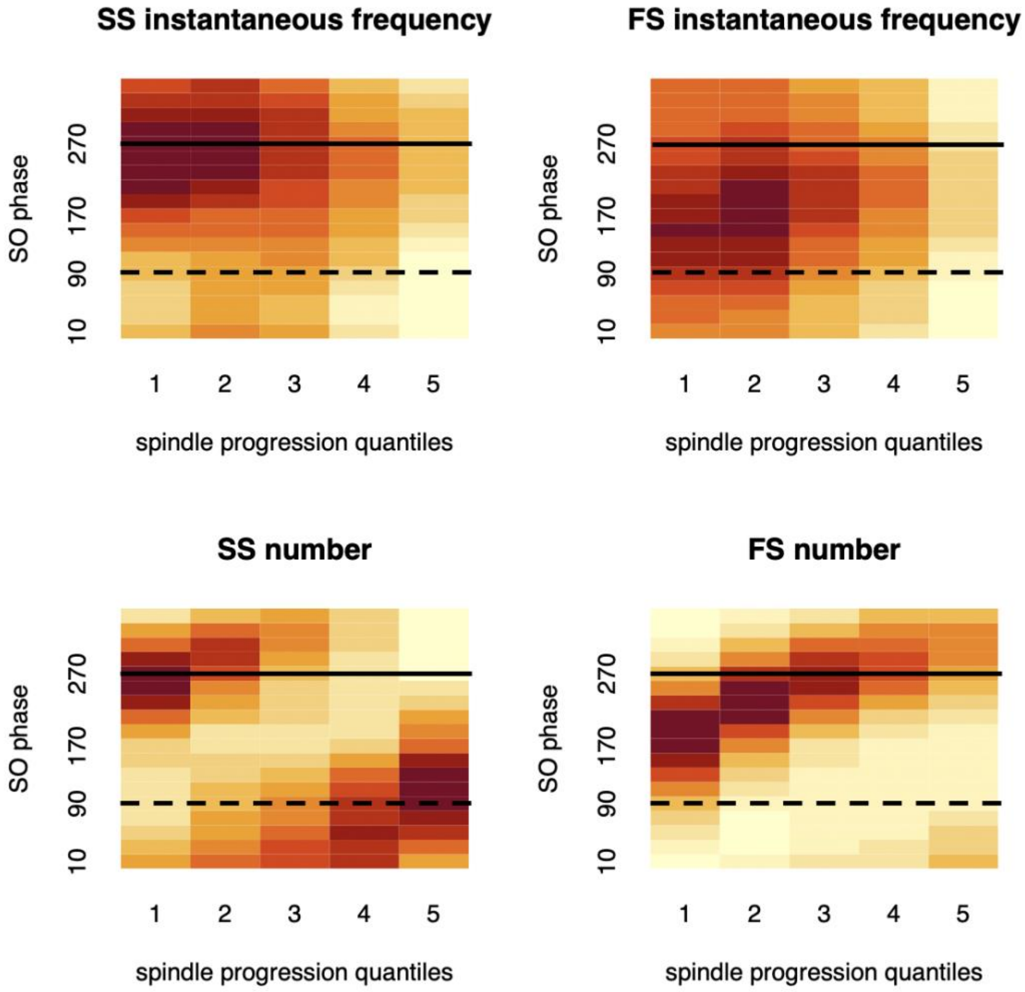


*Top two plots illustrate how slow spindle (SS) or fast spindle (FS) instantaneous frequency changes with respect to SO phase (vertical axis) and spindle progression (horizontal axis), averaged across all participants. The bottom plots similarly demonstrate how SS/FS count changes with respect to SO phase (i.e. reflecting conventional SO-spindle temporal coupling). The dashed horizontal black line (90 degrees) indicates the SO negative peak; the solid line (270 degrees) indicates SO positive peak.*

***Supplementary figure 3: Lack of strong association between wake and sleep EEG variables***

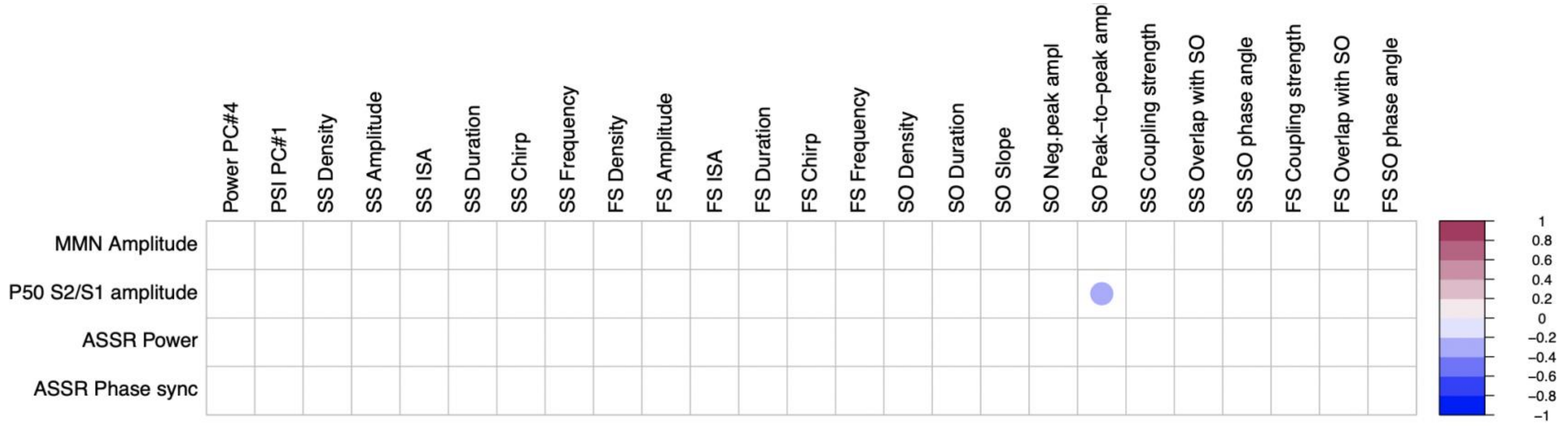


*Correlation matrix illustrating Pearson's correlations among 100 comparisons (4 wake by 25 sleep metrics at Cz) with abs(r)>0.2 and p<0.01. Correlations were computed based on all participant's data with effects of disorder, age and sex partialled out.*

*Supplementary figure 4: Associations between slow spindle density/amplitude and PANSS scores*

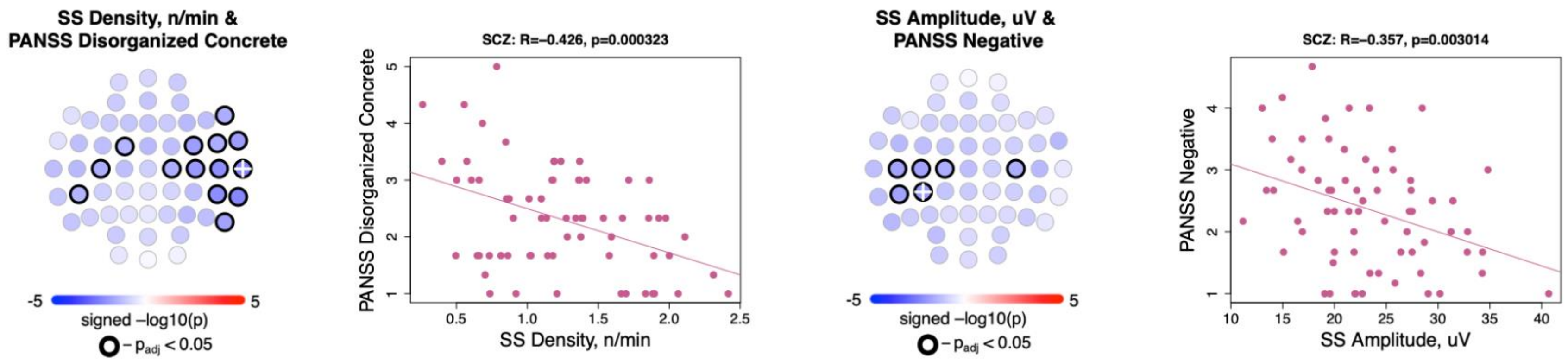


*Topoplots illustrate the significance of associations between SS density and PANSS Disorganized-concrete score (left); also, SS amplitude and PANSS Negative score (right). The scatterplots to the right of each topoplot illustrate SS density/amplitude as a function of PANSS score at EEG channel where association was the most significant (white cross mark on the topoplot). The title of each scatterplot also provides Pearson's correlation coefficient together with the p-value for this channel.*

***Supplementary figure 5: net PSI during N2 and REM in control subjects in GRINS and ESZ sample***

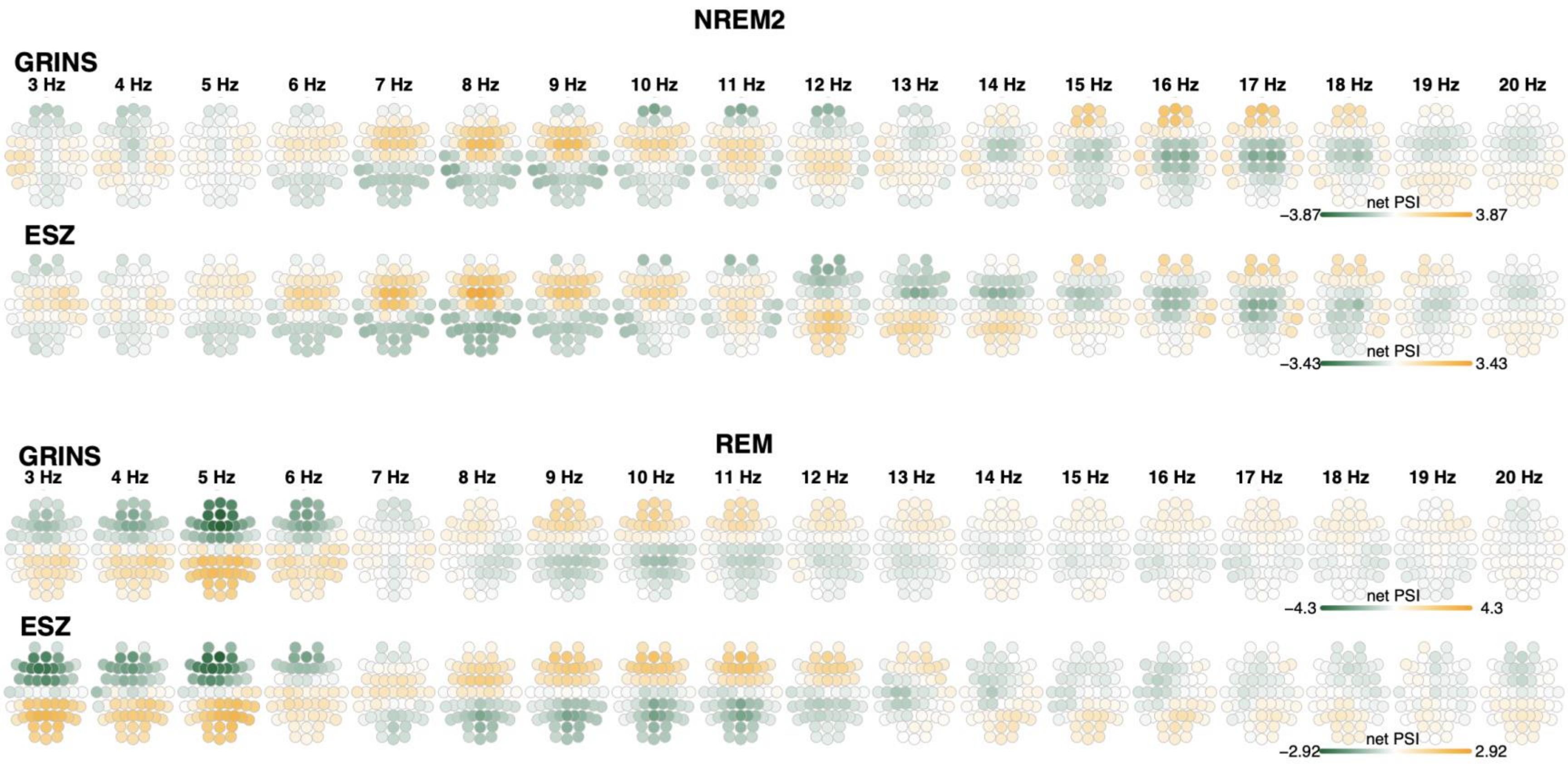


*The top two rows illustrate net PSI values during N2 for each frequency bin (center frequency varied from 3 to 20 Hz), with values averaged over the GRINS CTR group and the ESZ CTR group. For contrast, the bottom two rows illustrate the equivalent net PSI metrics during REM.*

### *Supplementary figure 6: PSD-based principal components used in SCZ/CTR prediction analysis*

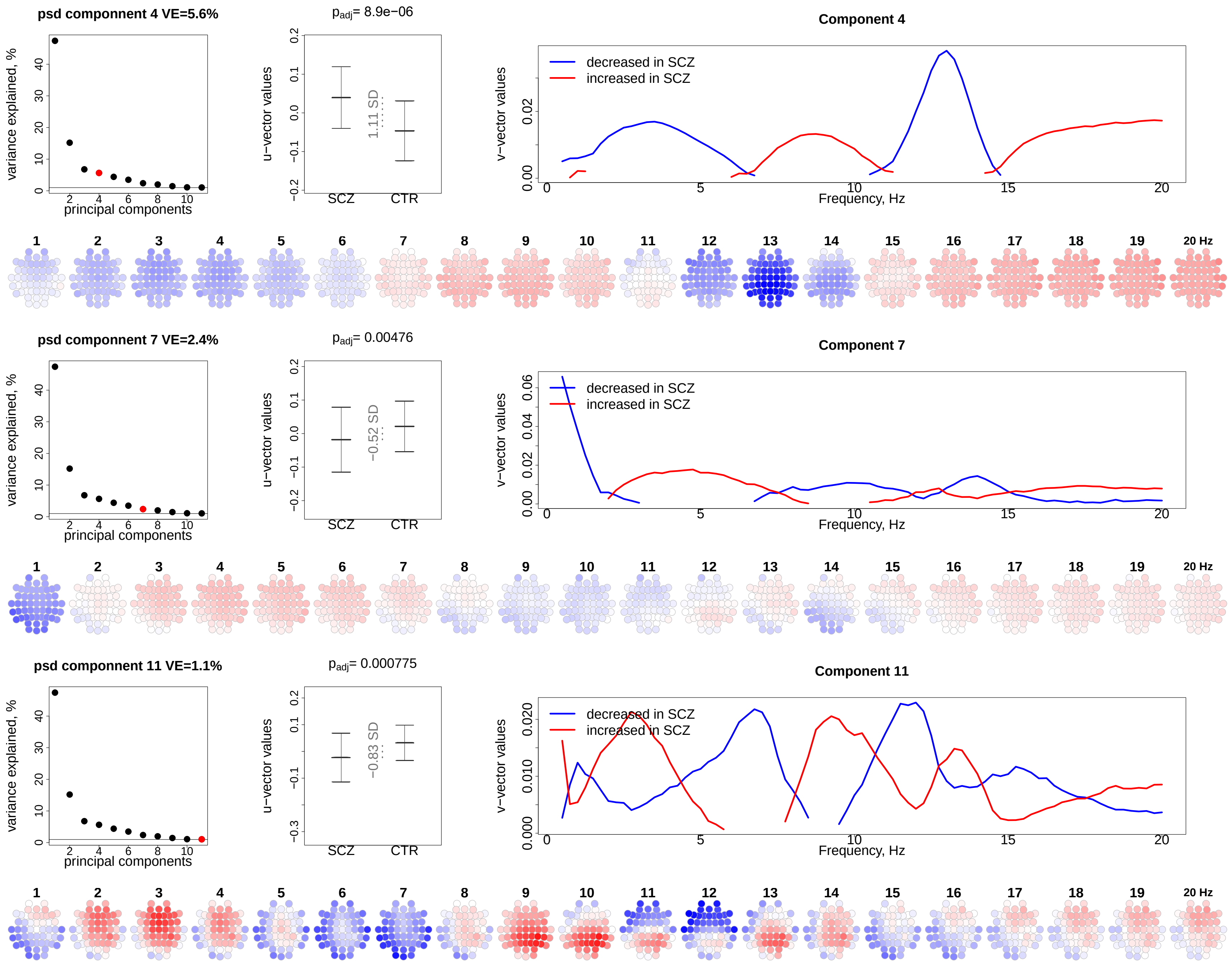

### *Supplementary figure 7: PSI-based principal components used in SCZ/CTR prediction analysis*

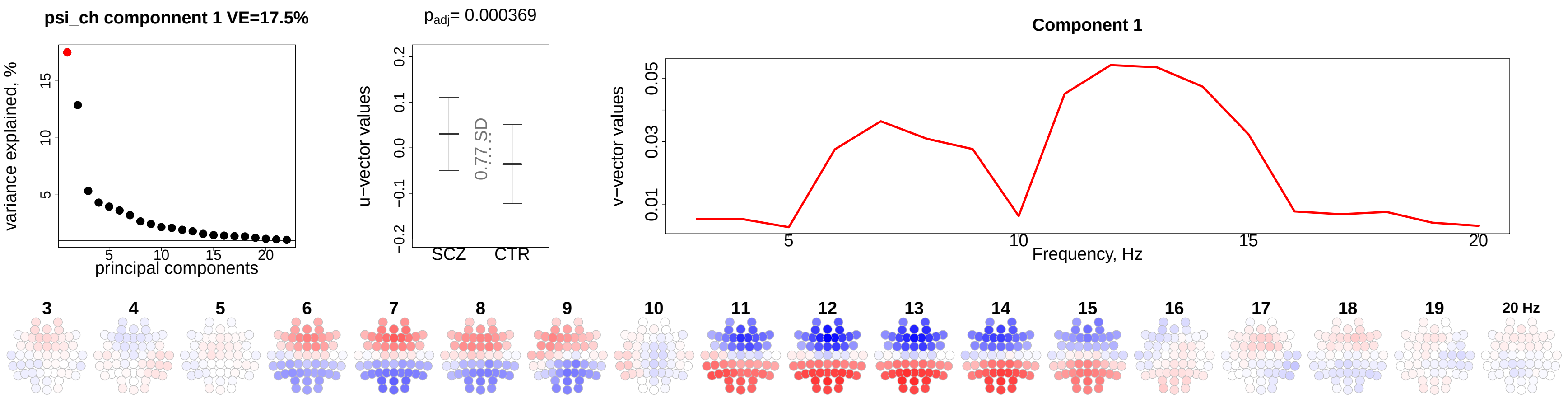


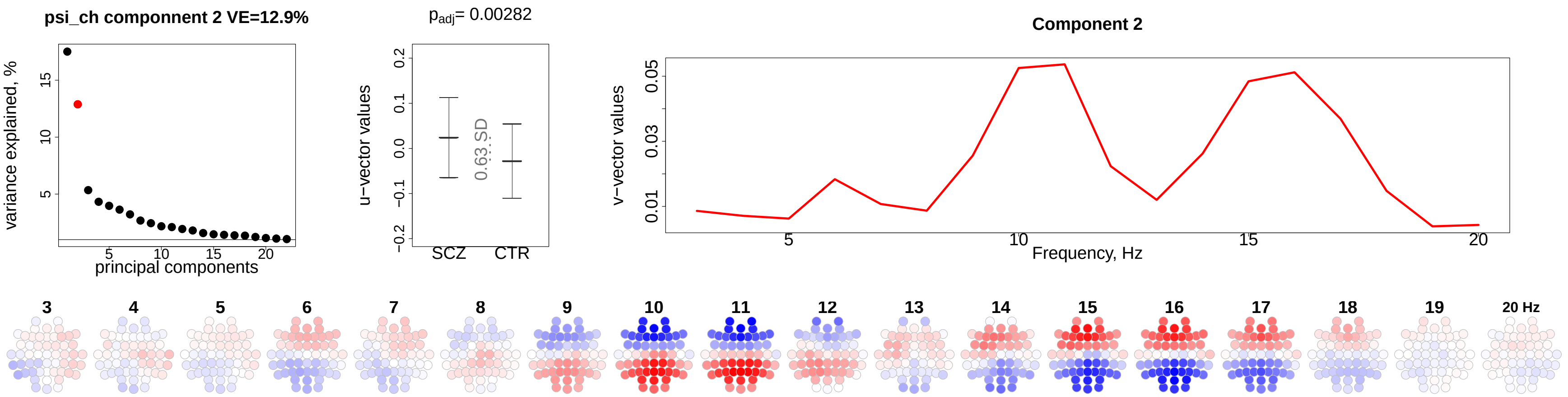

***Supplementary figure 8: Spindle-based principal components used in SCZ/CTR prediction analysis***

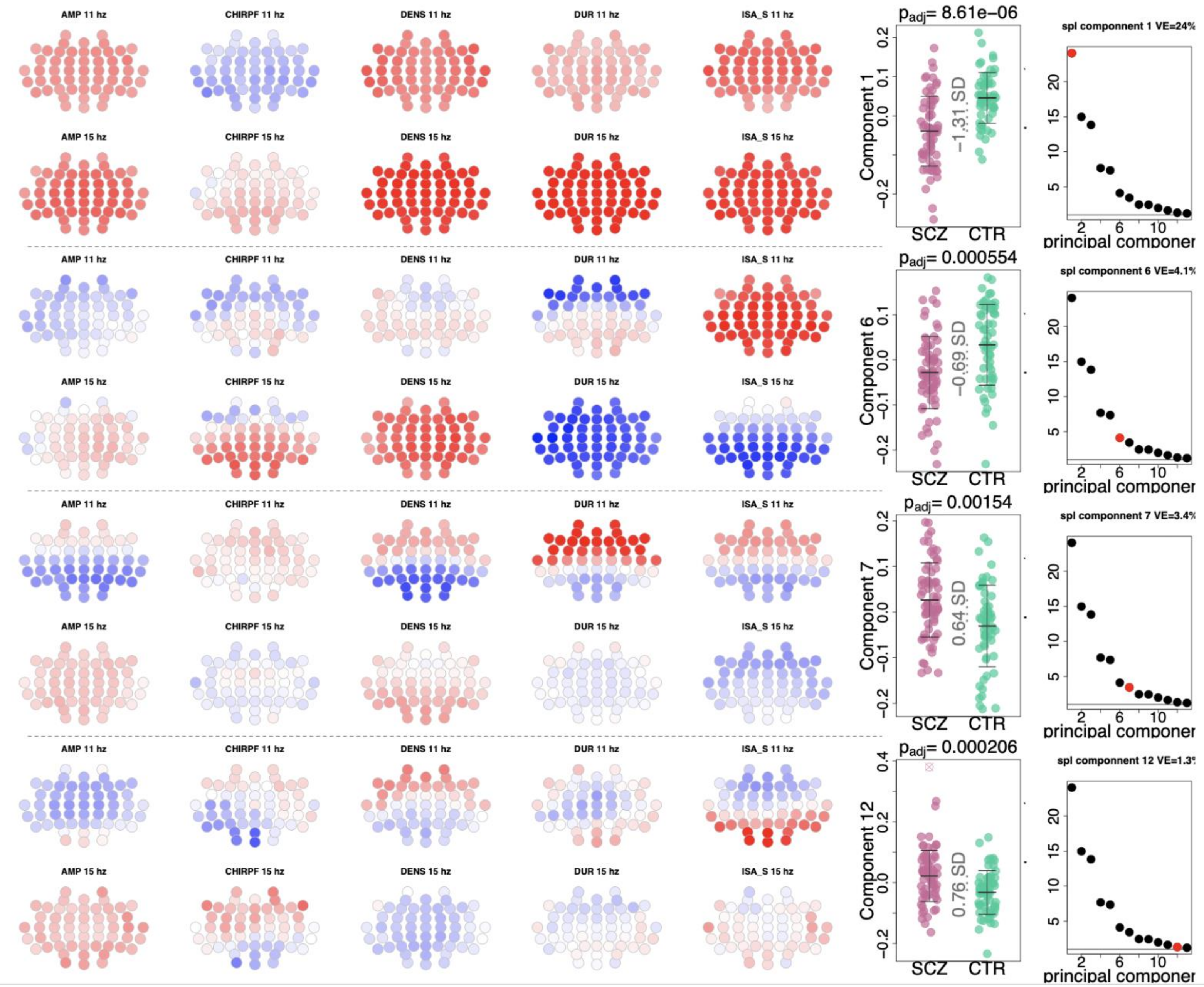

***Supplementary figure 9: SO-based principal components used in SCZ/CTR prediction analysis***

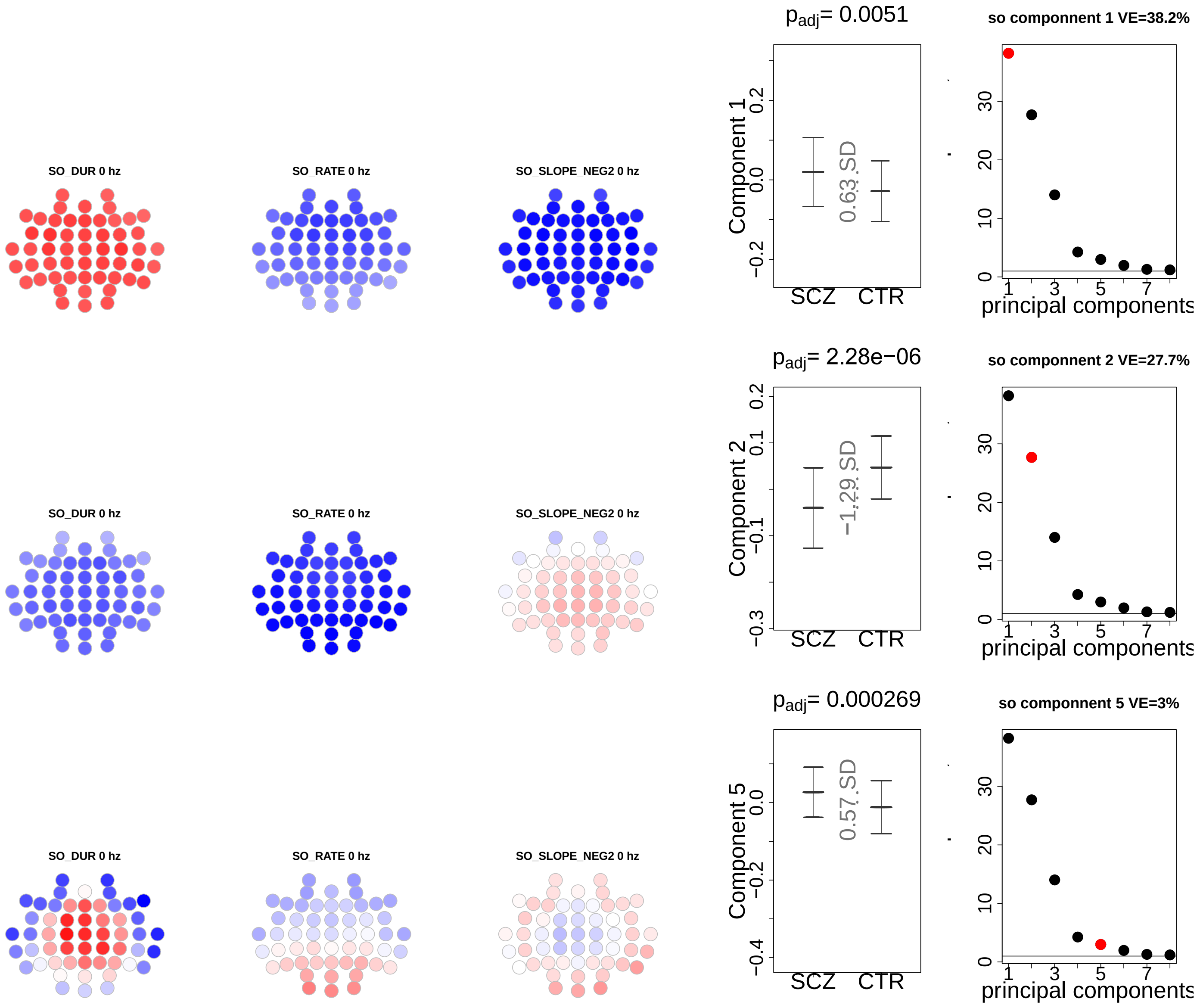

***Supplementary figure 10: SCZ/CTR prediction based on SS density at P7 and FS density at FC2***

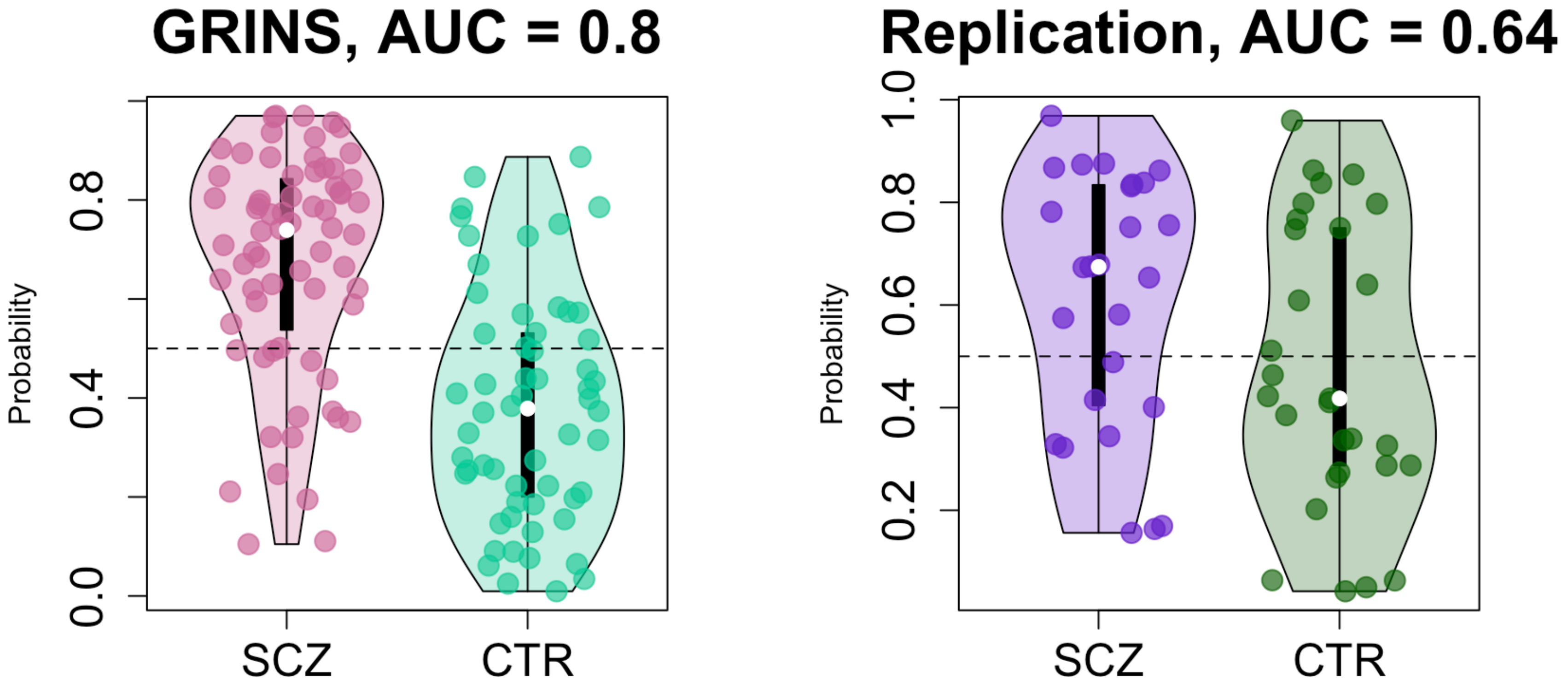


*Two violin plots the logistic regression probabilities for case/control labels for GRINS (training sample, left) and ESZ (independent target sample, right) using SS density at P7 and FS density at FC2 – channels with the largest effect sizes of group differences. AUC – area under the ROC curve.*
